# Orexin cells efficiently decode blood glucose dynamics to drive adaptive behavior

**DOI:** 10.1101/2022.04.14.488310

**Authors:** Paulius Viskaitis, Alexander L. Tesmer, Mahesh M. Karnani, Myrtha Arnold, Dane Donegan, Ed F. Bracey, Nikola Grujic, Daria Peleg-Raibstein, Denis Burdakov

**Affiliations:** Swiss Federal Institute of Technology (ETH Zürich), Department of Health Sciences and Technology, Zürich, Switzerland

## Abstract

Blood glucose variability shapes human brain performance and diverse clinical outcomes. However, it remains poorly understood how blood glucose fluctuations are decoded by genetically-defined neurons to change brain activity and behavior. Recent breakthroughs in genetics and clinical diagnostics identified hypothalamic hypocretin/orexin neurons (HONs) as core determinants of brain activity and adaptive behavior across mammals. Here we show that low-frequency HON population waves are tuned for transmitting information about minute-to-minute temporal features of blood glucose, thus rapidly converting its variability into brain state of behaving mice. Contrary to current theories envisioning glucose-proportional neural responses, the HONs’ response tracked blood glucose gradients, thus generating efficient neural adaptations in anticipation of maximal glucose deviations. Resolving this population response at the single cell level with volumetric multiphoton imaging furthermore revealed glucose-excited and glucose-inhibited HONs, distinctly coupled to body movements in the high-frequency domain. Finally, HON-selective opotogenetics and cell ablation demonstrated that HONs are critical for linking glucose to adaptive behavior. These results provide an insight into how behaviorally influential hypothalamic networks interpret blood glucose variability. This may inform future metrics for efficient prediction of glycemic states in health and disease.

## INTRODUCTION

Glucose is a fundamental vital variable in our bodies. It is released into the blood and taken up by tissues, resulting in fluctuations in circulating glucose levels on the time scale of minutes. These fluctuations predict and determine healthy and maladaptive physiological states (Gailliot and Baumeister, 2007; Gold, 1995; Messier and Gagnon, 1996; Zhou et al., 2020). Measuring and interpreting blood glucose variability has thus been a key focus of research in biology, medicine, and engineering (Campfield and Smith, 2003; Cobelli et al., 2011; Mergenthaler et al., 2013; Rodbard, 2018). Encoding of current glucose level provides information about available energy, however, for timely responses, it is also vitally important to predict future states. Future states can be inferred from the rate of change of a parameter (DiStefano et al., 2012). Therefore, the temporal derivative of blood glucose carries valuable information about future glycemic state (Cobelli et al., 2011; Marchetti et al., 2008). Indeed, reactive (feedback) and predictive (feedforward) control based on current states and derivatives of glucose improves performance of medical devices, such as the artificial pancreas (Borase et al., 2021; DiStefano et al., 2012; Marchetti et al., 2008). This raises an unanswered question: which features of blood glucose variability are sensed by the body’s own glucose-measuring systems, such as the brain’s glucose-sensing neurons (Bentsen et al., 2019; Burdakov et al., 2005; Routh, 2002)?

Here, we address this question at the level of a genetically-defined group of putative glucose-sensing neurons - the hypothalamic orexin/hypocretin neurons (HONs). These brain-wide projecting neurons promote wakefulness and orchestrate adaptive physiology across species (Sakurai, 2007). Their loss produces an inability to adapt arousal to body energy balance, as well as narcoleptic wakefulness instability and energy homeostasis disorders (Bassetti et al., 2019; Peyron et al., 1998; Sakurai et al., 1998; Yamanaka et al., 2003). Studies of glucose sensitivity in isolated HONs *in vitro* led to speculations that they may sense glucose *in vivo* (Burdakov, 2005; Gonzalez et al., 2008; Yamanaka et al., 2003). However, the sensitivity and physiological significance of HON glucosensing *in vivo* remain unknown because HON activity has never been correlated to measured changes in blood glucose, nor has the role of HONs in the body’s responses to glucose been described in behaving animals. Here, for the first time, we perform such measurements, and visualize blood glucose responses of the HON population, as well as of >900 individual HONs. Furthermore, by documenting how selective HON ablation or stimulation affects behavioral responses to glucose, we show the importance of HONs for these responses.

## RESULTS

### Temporal coupling of HON population signal to blood glucose

To correlate neural dynamics with concurrently measured changes in blood glucose concentration, we designed and implemented an experimental paradigm that combined telemetry of carotid electrochemical glucose sensors, and fiber photometry of HON-selective GCaMP6s fluorescent neural activity sensors (Figure 1A, B; see Methods).

**Figure 1.**
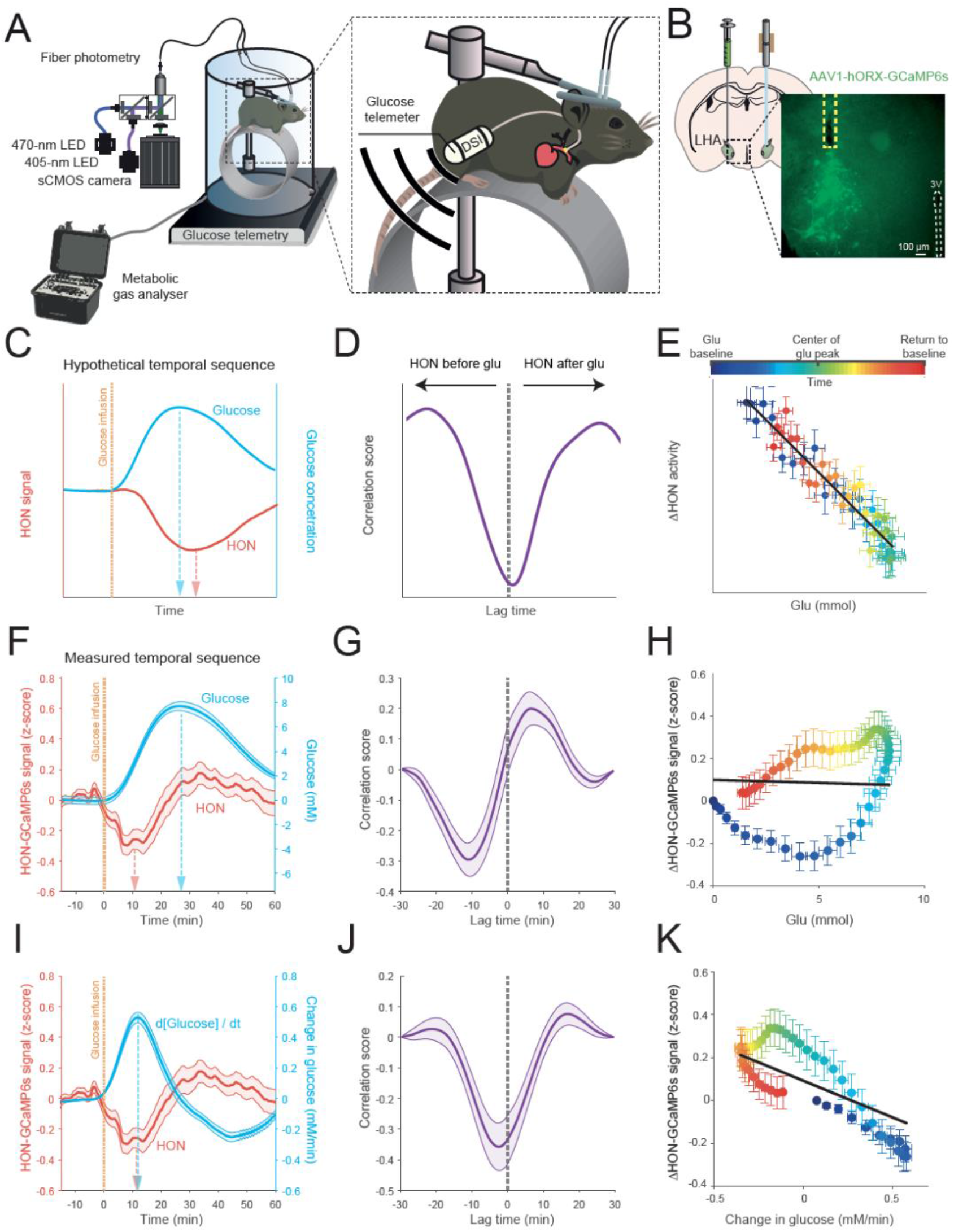
Temporal relations of HON population activity and blood glucose. **(A)** Scheme of experimental setup for simultaneous fiber photometry, indirect calorimetry, glucose and temperature telemetry (DSI), and locomotion recordings.**(B)** Stereotaxic surgery schematic (left), LHA lateral hypothalamic area, and (right) expression of GCaMP6s in HONs for fiber photometry. The dashed square box indicates the fiber location. 3V, 3rd ventricle. **(C-E)** Hypothetical temporal relations based on existing data (see Results text). **(F-H)** Measured temporal relations of HON activity and blood glucose, simultaneously recorded in the same experiments. **(F)** HON activity and blood glucose concentration traces; peak HON response preceded peak glucose concentration. Glucose infusion significantly decreased HON photometry signal (between 0 and 10 min) compared to baseline (−10 to 0 min), p < 0.0001, t = 4.146, df = 105. **(G)** Cross-correlation of HON activity and blood glucose. **(H)** Linear fit did not explain variability and fit slope was not different from zero: R2 = 0, p = 0.288. **(I-K)** Measured temporal relations of HON activity and glucose derivative (d[Glucose]/dt]). Linear fit explained some variability and the slope was significantly non-zero **(I)**: R2 = 0.04, p < 0.001. In C-K, data are means and SEM of n = 57 intragastric glucose infusion responses from 22 mice. Data smoothed by a 10-minute moving mean for visualization.

The current model of HON glucosensing predicts that HON inhibition should proportionally track absolute glucose concentration, with a sensing delay of a few minutes (Burdakov, 2005; Yamanaka et al., 2003). If such concentration-proportional sensing happened in the intact organism whose blood and brain glucose levels rapidly equilibrate (Silver and Erecinska, 1994), we would expect maximal HON inhibition to occur after glucose peak (Figure 1C, D), and a negative monotonic relation between HON activity and increasing glucose concentration (Figure 1E).

However, we found that *in vivo*, a rise in blood glucose rapidly and significantly inhibited HON population activity (Figure 1F), regardless of the route of glucose delivery (intragastric or intraperitoneal; Figure S1). However, contrary to the current model, the timing of HON inhibition strikingly diverged from a time-delayed copy of the blood glucose waveform. Instead, the HON inhibitory response was seen only during blood glucose rise, with peak HON response occurring several minutes before – rather than after – the blood glucose peak (Figure 1F, G). Also unexpectedly, HON activity subsequently returned to control levels when glucose was at peak level and stable, and then rose when glucose was falling (Figure 1F, G). As a result, HON activity state as a function of blood glucose displayed a hysteresis profile (Figure 1H), rather than the expected linear relationship (Figure 1E). From this noncanonical temporal profile of the HON glucose response, we hypothesized that the HON inhibition tracked the rate of change of glucose (i.e. its first derivative, d[glucose]/dt). Indeed, re-plotting differentiated glucose data revealed that HON inhibition is a close mirror-image of the glucose derivative (Figure 1I, J), and there is a significantly negative linear relationship between HON and the glucose derivative (Figure 1K). This relationship was similar irrespective of route of glucose administration (intragastric: Figure 1K; interaperetoneal: Supplementary Figure 1)

Together, these data show that HON population is rapidly inhibited by a rise in blood glucose, and this inhibition tracks the rate of change, and not absolute values, of blood glucose.

### HON population dynamics preferentially encode blood glucose dynamics

Changes in blood glucose level are expected to affect multiple behavioral and metabolic parameters, and HON activity may co-vary with a number of those (Karnani et al., 2020; Williams et al., 2007). This raises a fundamental and currently unanswered question: do HONs specialize in glucose sensing, or is glucose influence relatively minor compared to the other variables that may affect HONs *in vivo*?

To answer this, we needed simultaneous experimental tracking of multiple physiological variables, and a method to quantify how much HON activity variability is attributed to each separate variable, despite the presence of multiple variables. To this end, we designed an experimental and analysis workflow enabling the co-monitoring of multiple variables at the same temporal resolution, including blood glucose, CO_2_ and O_2_ respiratory gas exchange, HON-GCaMP6s activity, body temperature, and locomotion (Figure 1A, Figure 2A). We then used the resulting data matrix (Figure 2A-G) to quantitatively predict the HON population activity based on the other physiological variables (“predictors”) with an encoding model (Engelhard et al., 2019) (Figure 2H). Predictors also included the first derivatives of variables (Figure 2H), because this temporal feature was implicated in HON responses to glucose (Figure 1I-K).

**Figure 2.**
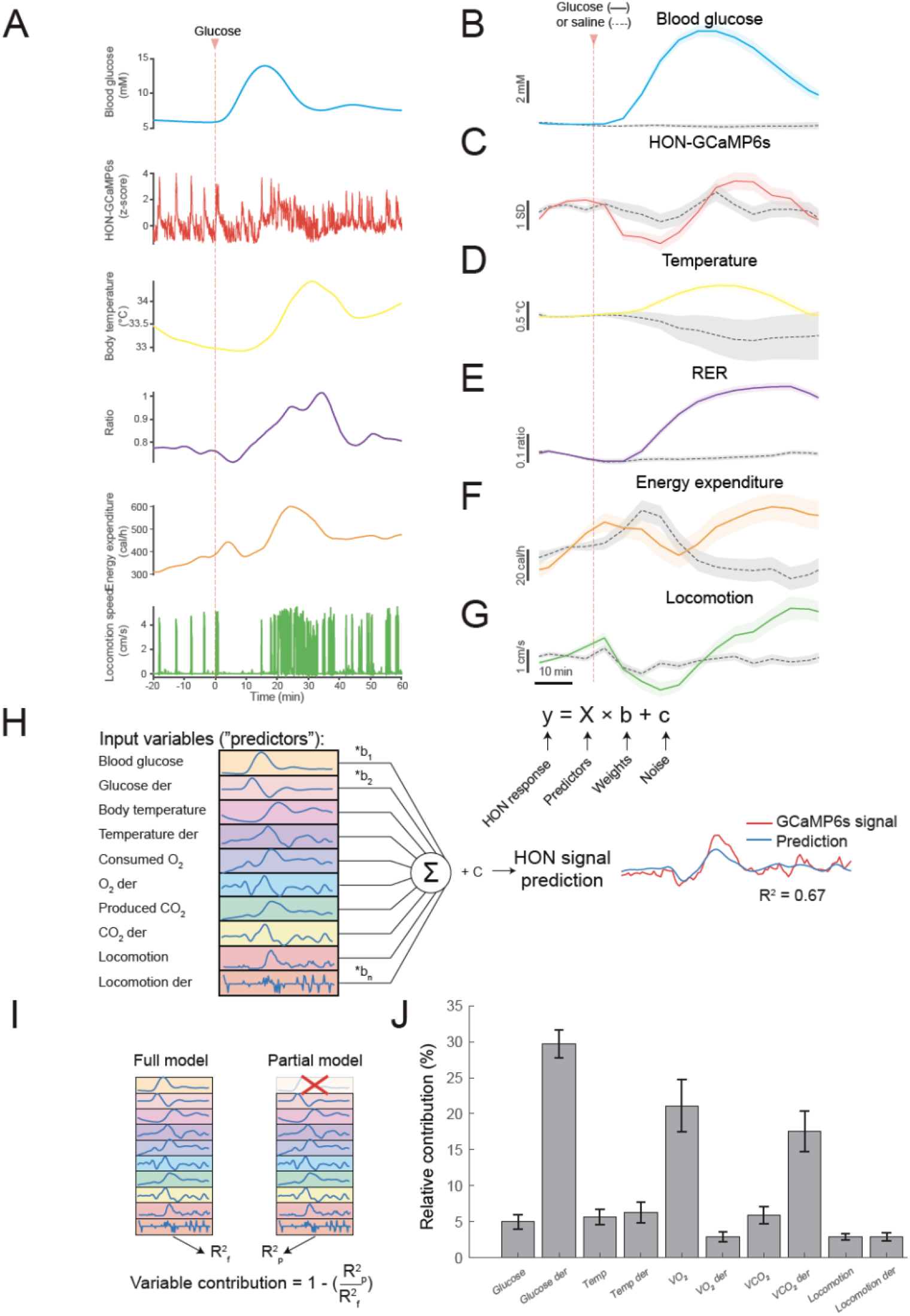
Relative influence of behavioral and metabolic variables on HONs. **(A)** Example multi-parameter data from a single recording session. **(B-G)** Group data for the experiment show in A. Data shown in 5 min bins for visualization only. The 5 – 25 minutes epoch was used for statistics. Compared to saline infusion, glucose injection: **B**, increased blood glucose level (t-test p < 0.0001; df = 26, tstat = −5.563; n = 7 & 21 responses from 6 and 6 animals to sal and glu respectively), **C**, reduced HON activity responses (t-test p < 0.001; df = 69, tstat = 3.918; n = 39 & 32 responses from 13 and 10 animals to sal and glu respectively), **D**, increased body temperature (t-test p < 0.05; df = 31, tstat = −2.598; n = 7 & 26 responses from 7 and 8 animals to sal and glu respectively), **E**, increased respiratory exchange ratio (RER) (t-test p < 0.0001; df = 87, tstat = −6.322; n = 59 & 30 responses from 30 and 15 animals to sal and glu respectively), **F,** did not alter energy expenditure (t-test p = 0.08; df = 87, tstat = 1.762; n = 59 & 30 responses from 30 and 15 animals to sal and glu respectively) and **G,** reduced running (t-test p < 0.0001; df = 117, tstat = 4.736; n = 80 & 39 responses from 32 and 19 animals to sal and glu respectively). **(H)** Depiction of the general linear model workflow to predict HON activity from the simultaneously-measured set of physiological variables. **(I)** Illustration of input variable contribution testing. **(J)** Ranking of input variable contributions for accuracy in modeling HON activity after glucose infusion. Data from 17 recording sessions from 5 mice. 10-minute time bar shown in panel I also applies to B-G. Data are means and SEM.

Using this encoding model, we calculated the relative contribution of each physiological variable to the slow HON population response, by quantifying how much of the explained variance decreased when that variable was removed from the model (Figure 2I). The highest relative contribution was attributed to the derivative of glucose (29.7±2% relative contribution to explained variance), followed by consumed oxygen volume, (VO_2_, 21.1±3.6%), and by derivative of produced carbon dioxide volume (VCO_2_, 17.6±2.8%) (Figure 2J; confirmed for a variety of glucose infusion parameters in Supplementary Figure 2).

Thus, when co-variation of HON activity across multiple physicochemical and behavioral factors is considered, the glucose derivative emerges as a strong determinant of information transmitted by the HON population.

### Individual HONs report diverse temporal features of blood glucose

We next sought to determine whether blood glucose homogeneously affects individual HONs. This is important because it is not clear how individual neurons contribute to the overall population signal measured by the photometry. Furthermore, it has been recently demonstrated that photometry signal may sometimes originate from nonsomatic changes in calcium (Legaria et al., 2022). We therefore switched our HON activity recording mode from bulk photometry to single-cell resolution using 2-photon volumetric imaging of HONs through hypothalamus-implanted gradient index (GRIN) lenses (Karnani et al., 2020) (Figure 3A). We confirmed that, at population level, HON glucose responses were similar across the two recording modes, by observing comparable glucose responses in fiber photometry and summed 2P HONs imaging (Figure 3B-D).

**Figure 3.**
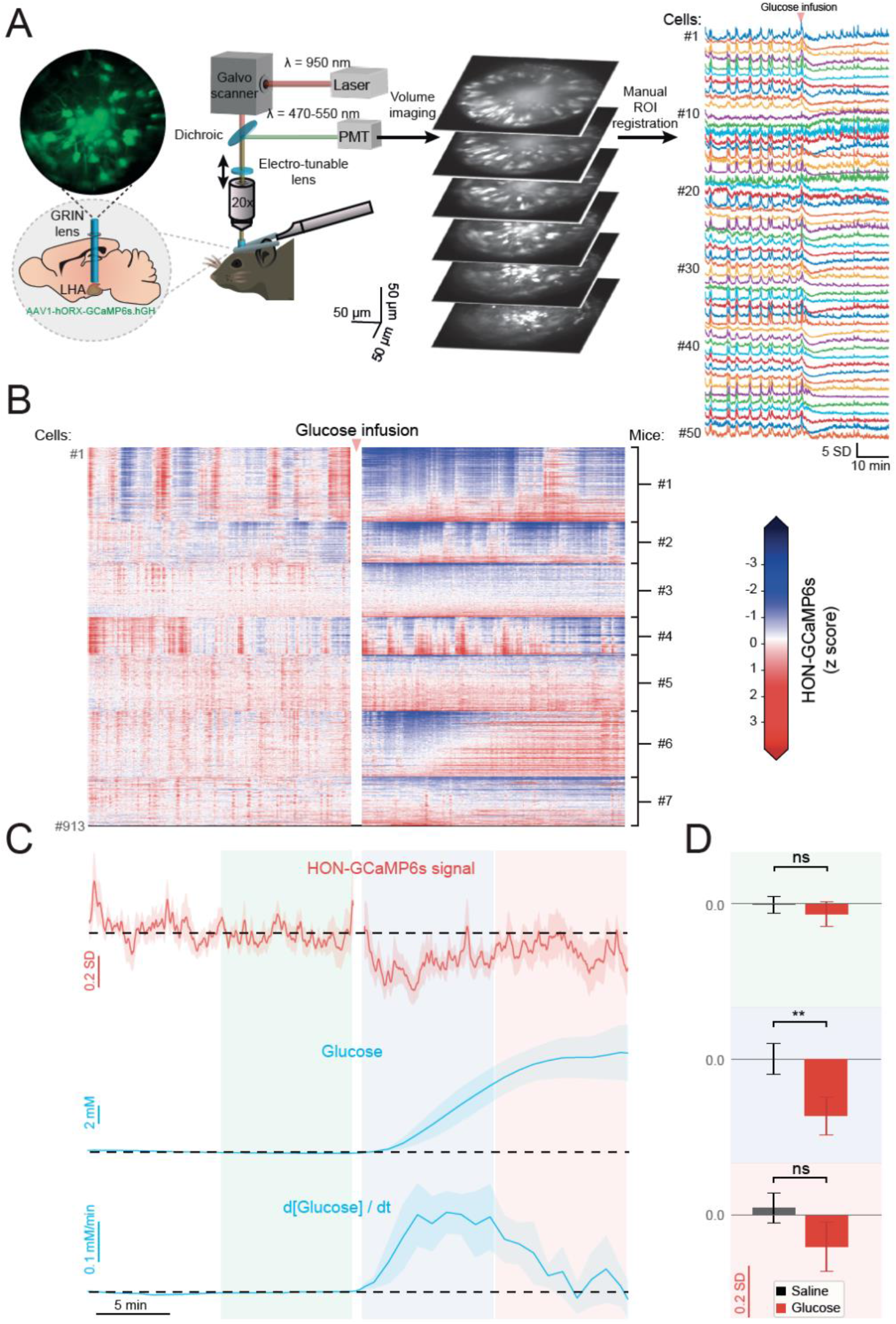
Single-cell resolution analysis of HON responses to blood glucose fluctuations. **(A)** Scheme of 2-photon imaging of individual HONs in behaving mice. **(B)** Individual responses of 913 HONs to glucose infusion from 7 mice (saline control is shown in Supplementary Figure 3). **(C)** Temporal alignment of average HON responses (top) to blood glucose concentration (middle) and its derivative (bottom). **(D)** HONs were significantly inhibited only during high glucose derivative phase (2 to 11 min, blue box), but not during high absolute glucose (11 to 20 min, red box) or during baseline periods (−11 to −2 min, green box). 2-way repeated measures ANOVA: treatment p = 0.029, F (1.0, 6.0) = 8.216; Šidák correction multiple comparisons: p = 0.709 (−11:−2), p = 0.005, (2:11), p = 0.534 (11:20); n = 7 mice. Breaks in the neural activity recordings are due to laser shutter being closed during infusions in majority of the experiments. Data are presented as means and SEM; **p < 0.01 and ns p > 0.05.

In each mouse (n = 7), we simultaneously resolved the activity of 92-180 HONs, analyzing a total of 913 HONs from 7 mice (the full dataset is shown in Figures 3B and Supplementary 3). To compare glucose responses of individual cells, we fitted each cell’s activity profile to several templates representing distinct temporal features of glucose dynamics, and classified cells based on best goodness-of-fit (Figure 4A, see Methods). Based on this classification, the majority of HONs (98%; 895/913 cells) responded to blood glucose dynamics, with only 2% of cells not fitting any of the templates (Figure 4B, D). The majority of the glucose-responding HONs were glucose-inhibited (64%; of which 31% were glucose-derivative-inhibited cells, and 33% were glucose-proportional-inhibited cells), but we also detected other functional subclasses (glucose-derivative-activated cells, 11%, glucose-proportional-activated cells, 25%). (Figure 4B, D). The summed/population level response (Figure 3C, Figure 1) resembled that of glucose-derivative-inhibited cells (idG cells, Figure 4B), presumably because of the small response amplitudes of glucose-derivative-activated cells (dG, Figure 4B), and relatively similar response amplitudes of glucose-proportional-inhibited and glucose-proportional-activated cells (iG and G cells respectively, Figure 4B). We could also recreate the HON population response to glucose infusion by multiplying the glucose templates by the relative HON-subgroup prevalence (Supplementary Figure 4A). The HON subsets were largely intermingled within the recording volume (Figure 4E).

**Figure 4.**
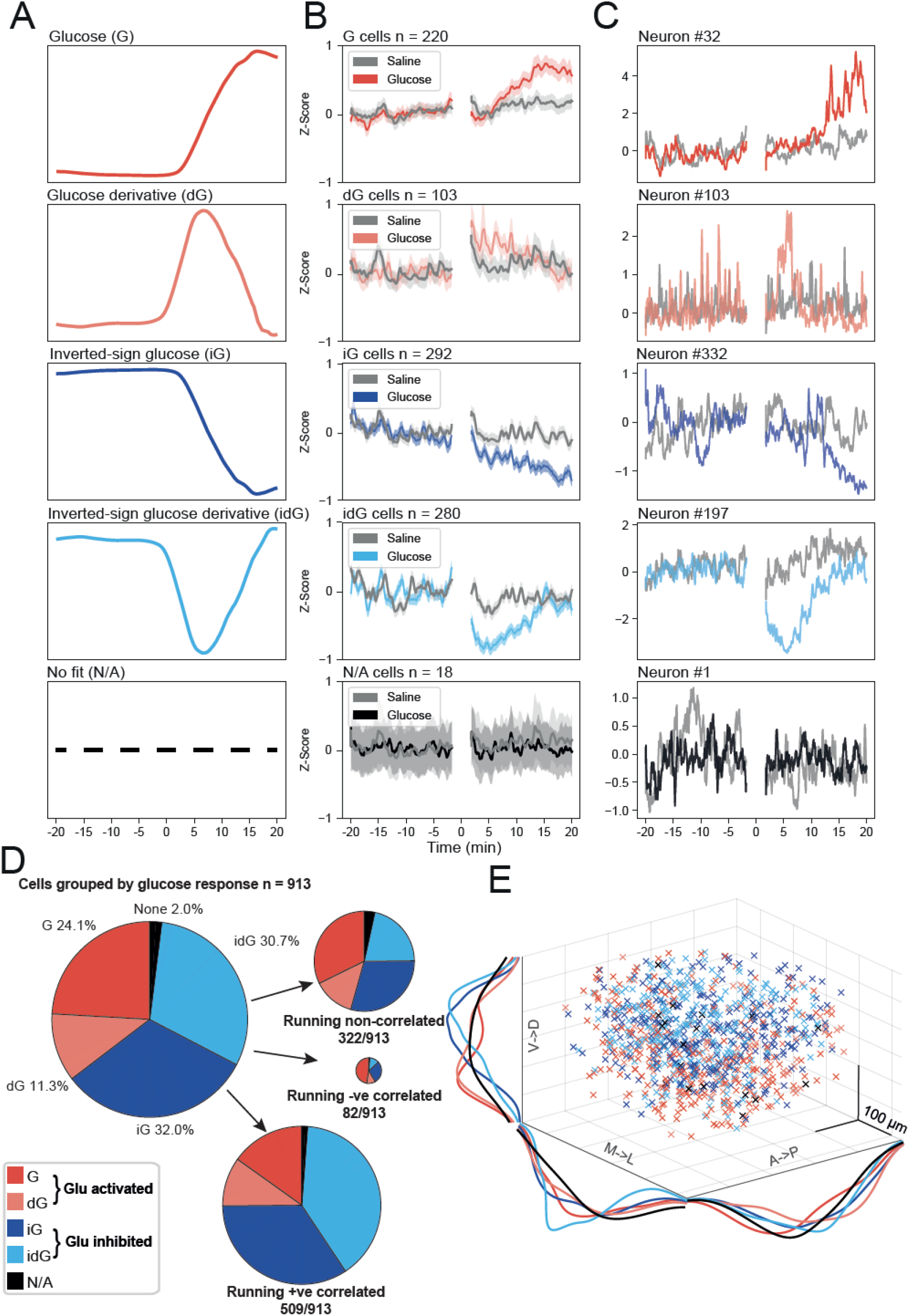
Classification of single-cell HON responses to blood glucose and running. **(A)** Fitting templates based on recorded blood glucose dynamics: glucose (G), glucose derivative (dG), inverted glucose (iG), inverted glucose derivative (idG) and no fit (N/A). **(B)** Average HON activity traces for each of the glucose-response classes. **(C)** Examples of individual neuron from each class. **(D)** Relative proportions of HON subsets with regards to their responses to glucose and further subdivided by correlations with running. **(E)** Anatomical distribution of HON subsets. Breaks in the neural recordings are due to laser shutter being closed during infusions in majority of the experiments. Data are presented as means and SEM.

We next asked how other information is distributed across these glucose-classified HONs. On rapid timescales, individual HONs display heterogeneous temporal coupling to locomotion, with some neurons positively correlated, some negatively correlated, and some noncorrelated, with running (Karnani et al., 2020). We re-examined our new data from this perspective of running, which we recorded concurrently with neural activity in all 2P-GRIN experiments. We found that the majority of running-correlated HONs were glucose-inhibited while the majority of running-negatively-correlated HONs were glucose-excited (Figure 4D, right). In agreement with this, glucose inhibited cell subpopulations had larger proportions of cells positively correlated with running than the other classes of HONs (Figure S4B).

Overall, these data indicate that, while there are distinct subpopulations of HONs in terms of correlations with blood glucose dynamics or running activity, the bulk of HONs are glucose-inhibited (iG and idG cells in Figure 4B), and the majority of individual HONs are positively correlated with running and glucose-inhibited.

### Role of HONs in behavioral responses to glucose

Blood glucose governs the function of blood glucose regulators in the hypothalamus and pancreas (Gerich et al., 1976; Pozo and Claret, 2018), and may also govern behavior. We considered the possibility that the inhibition of bulk HON activity by glucose could be involved in these processes. If this HON response to glucose is involved in blood glucose control, then glucose dynamics may be altered in HON-ablated animals. To test this, we selectively ablated HONs in adult mice using the orexin-DTR mouse model (Gonzalez et al., 2016b) (Figure 5A). Glucose-tolerance tests, however, did not reveal any major differences in glucose dynamics between control mice and HON-ablated mice (Supplementary Figure 5A, B).

**Figure 5.**
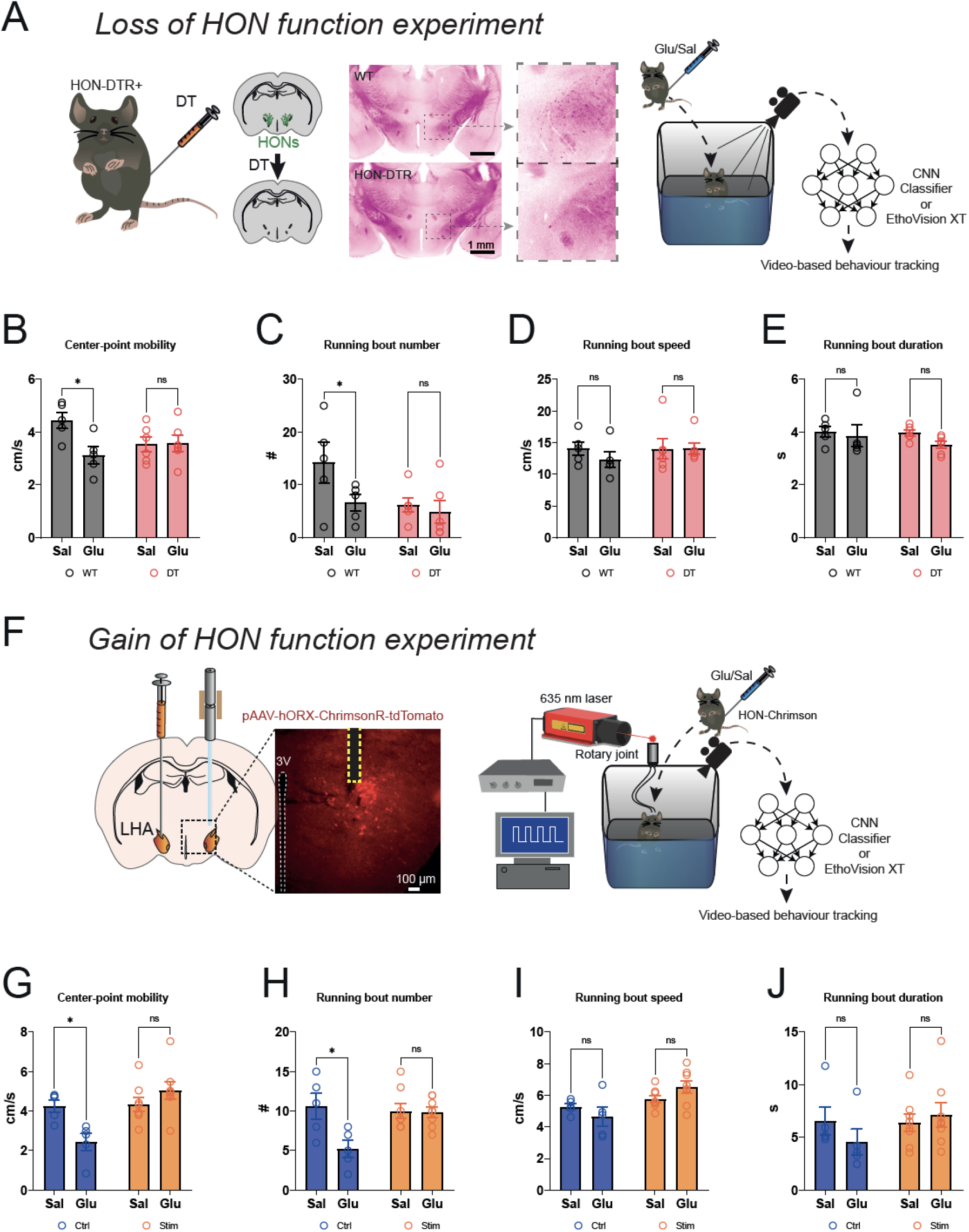
Role of HONs in glucose-evoked behavior. **(A)** Left, scheme for selective HON ablation strategy; middle, example histology from HON-ablated HON-DTR (DT) mice or wild-type (WT) control and right, glucose-induced behavior assessment study design diagram. **(B)** Mean running speed during the 10 minutes of open field experiment in saline (sal) or glucose (glu) injected WT and DT mice. Glucose significantly suppresses running speed in WT but not in DT mice. 2-way repeated measures ANOVA: treatment x group F (1, 9) = 5.4, p = 0.04; WT sal vs glu p = 0.03; DT sal vs glu p > 0.99. **(C)** Number of machine-learning identified running bouts from experiment in A. Glucose significantly lowered number of running bouts in WT but not in DT mice. 2-way repeated measures ANOVA: treatment x group F (1, 9) = 5.3, p = 0.04; WT sal vs glu p = 0.04; DT sal vs glu p = 0.89. **(D)** Mean running speed during running-bouts. Glucose did not significantly alter this parameter in WT or DT mice. 2-way repeated measures ANOVA: treatment x group F (1, 9) = 0.71, p = 0.42. **(E)** Duration of running bouts was not significantly changed in WT or DT mice. 2-way repeated measures ANOVA: treatment x group F (1, 9) = 0.71, p = 0.42. **(F)** Selective HON activation strategy; histology HON-Chrimson mouse brain; and glucose-induced behavior assessment study design diagram. **(G)** Mean running speed during the 10 minutes of open field experiment from **F** in saline (sal) or glucose (glu) injected Control (Ctrl) and optogenetic stimulated HON-Chrimson (Stim) mice. Glucose significantly suppresses running speed in Ctrl but not in Stim mice. 2-way repeated measures ANOVA: treatment x group F (1, 11) = 13.78, p < 0.01; Ctrl sal vs glu p = 0.01; Stim sal vs glu p = 0.23. **(H)** Number of machine-learning identified running bouts from experiment in F. Glucose significantly lowered number of running bouts in Ctrl but not in Stim mice. 2-way repeated measures ANOVA: treatment x group F (1, 11) = 19.11, p < 0.01; Ctrl sal vs glu p < 0.01; Stim sal vs glu p = 0.46. **(I)** Mean running speed during running-bouts. Glucose did not significantly alter this parameter in Ctrl or Stim mice. 2-way repeated measures ANOVA: treatment x group F (1, 11) = 3.57, p = 0.09. **(J)** Duration of running bouts was not significantly changed in Ctrl or Stim mice. 2-way repeated measures ANOVA: treatment x group F (1, 11) = 1.45, p = 0.25. Data are presented as means and SEM; *p < 0.05, **p < 0.01, and ns p > 0.05.

We next examined the idea that glucose inhibition of HONs shapes behavior. Of note, running is stimulated in proportion to bulk HON activation (Karnani et al., 2020). Thus, if HONs are a critical determinant of glucose-modulated movement, the following predictions can be made. First, glucose infusions should suppress movement in the same time-window where bulk HON inhibition is seen (i.e. 0-10 min, see Figure S1A, 4C). Second, selective HON ablation should be sufficient to produce a similar movement suppression. Third, glucose infusions in HON-ablated mice should produce no further suppression of movement. Fourth, selective HON activation should prevent glucose from suppressing movement.

To test these predictions, we compared movement baselines and responses to glucose in HON ablated mice (“loss of function experiment”, Figure 5A-E) and in mice whose HONs were kept in a high activity state by HON-targeted tonic optogenetic stimulation (“gain of function” experiment, Figure 5F-J). Our data were in line with above predictions. First, glucose infusions suppressed movement (WT mice in Figure 5B, Ctrl mice in Figure 5G). This was associated with reduced number of running bouts but no change in bout speed or duration (WT mice in Figure 5C-E, Ctrl mice in Figure 5H-J). Second, movement suppression was induced by HON ablation ((Karnani et al., 2020), and reproduced here in Figure 5B-E). Third, glucose infusions in HON-ablated mice did not produce further movement suppression (Figure 5B-E, DT mice). Fourth, when glucose infusions were performed on the background of selective optogenetic HON activation, glucose did not suppress movement (Figure 5G-J).

Together, these loss and gain of function results thus indicate that selective HON manipulations (both gain and loss of function) are sufficient to block normal behavioral responses to glucose.

## DISCUSSION

Blood glucose variability shapes human brain performance and diverse clinical outcomes (Gailliot and Baumeister, 2007; Gold, 1995; Messier and Gagnon, 1996; Zhou et al., 2020). However, it remains poorly understood how blood glucose, brain activity, and behavior are linked at the level of genetically-defined neurons. Recent breakthroughs in genetics and clinical diagnostics identified HONs as core determinants of brain activity in both mice and humans (Chemelli et al., 1999; Hara et al., 2001; Nishino et al., 2000; Nishino et al., 2001). Here we show HONs rapidly respond to blood glucose fluctuations *in vivo*, thus tightly linking blood glucose variability and brain activity in behaving animals. Furthermore, by HON-selective loss and gain of function experiments, we find that HONs are critical for linking glucose to behavior. These results provide an insight into how behaviorally influential hypothalamic networks interpret blood glucose variability.

### Temporal multiplexing of blood glucose variability and behavioral state in HONs

The unprecedented simultaneous real-time measurements of blood glucose and HON dynamics reported here reveal unforeseen complexity of HON activity patterns inside the living organism. Concurrently with the realization that HONs t rapidly alter vital behaviors (Giardino et al., 2018; Karnani et al., 2020; Sakurai, 2014; Viskaitis et al., 2022), much work focused on external stimuli controlling their activity *in vivo*. This revealed that HON activity is acutely regulated by diverse external stimuli at the time scale of milliseconds (Gonzalez et al., 2016a; Hassani et al., 2016; Karnani et al., 2020; Mileykovskiy et al., 2005). Our new data show that HONs are also specialized to encode fluctuations in blood glucose at a slow (minute) time-scale (Figure 2). At the population level, HONs represented blood glucose trends (temporal derivative), rather than absolute levels (Figure 1). Within this population, however, some HONs responded proportionally to absolute level of blood glucose (Figure 4).

A key finding of the present study is that information about blood glucose and running is integrated within the same individual HONs (Figure 4D). This ability of individual HONs can be viewed as a form of temporal multiplexing, or distributed coding, i.e. carrying multiple streams of information at the same time in the form of a single, complex signal. This provides an important insight into organization of HONs, because it was previously unclear whether, *in vivo*, locomotion and glucose are encoded within the same HONs.

To the best of our knowledge, it has not been previously established whether the brain’s glucose-sensors transmit information about actual blood glucose level and/or its temporal trends. The latter question is fundamental for any sensor (Carpenter and Reddi, 2012; DiStefano et al., 2012), and of specific importance for understanding of blood glucose sensors, and their clinical replacement with closed-loop insulin pumps (Cobelli et al., 2011; Marchetti et al., 2008; Pinsker et al., 2016). The ability of HONs to communicate blood glucose derivatives (Figure 1,2), as well as display diverse responses at the single cell level (Figure 4), is surprising. It implies that HONs are not simply directly inhibited by glucose as previously predicted (Burdakov et al., 2005), but integrate additional upstream information – likely from other glucose-sensing endocrine or neural systems - resulting in the observed *in vivo* glucose responses. The nature of these upstream modulations that ultimately shape the blood glucose interpretation of HONs is as yet unknown, and will be a key subject for future investigations. Overall, our findings thus reveal a fundamental logic of how blood glucose sensors operate in the brain.

### HONs as a causal link between glucose and behavior

Our behavioral analyses (Figure 5) indicate that HONs do not merely track blood glucose fluctuations, but participate in using this information to adjust body movements. After glucose injections, running was rapidly and significantly decreased relative to control saline injections (Figure 5). Importantly, this behavioral effect occurred during the time interval associated with blood glucose rise and HON inhibition (Figure 1), and was abolished by optogenetic “activity clamping” of HONs (Figure 5F-J), or by HON ablation (Figure 5A-E). Together, these observations indicate that HONs are essential for glucose-induced control of running. It is tempting to speculate that this evolved to prevent moving away from food (Viskaitis et al., 2022), and/or to reduce movement-related energy use, when blood glucose is rising thus facilitating body energy repletion. From this perspective, it is interesting that glucose-induced running suppression can be overridden by selective HON activation (Figure 5F-J), because physiologically HONs are activated by diverse salient signals, such as those indicative of stress or reward (Gonzalez et al., 2016a; Hassani et al., 2016; Karnani et al., 2020; Peleg-Raibstein and Burdakov, 2021). In future work, it would therefore be important to test whether glucose-induced suppression of locomotion can be overridden by salient signals that activate HONs, which might provide an evolutionary advantage by permitting behavioral flexibility.

### Implications for understanding the brain’s strategies for efficient anticipatory control

Understanding the general logic of brain function is a long-established goal in biology, especially relevant today due to the growing cross-talk between neuroscience and engineering (Hassabis et al., 2017; Richards et al., 2022; Schiff, 2011). It therefore seems appropriate to conclude by speculating about implications of our findings from an engineering perspective. A fundamental shared concept in neuroscience and control engineering is that of anticipatory (aka feedforward, predictive) control. This refers to the ability to prepare for predictable future events, which constitutes a clear survival advantage. An elementary algorithm for anticipatory responses is “derivative control”, where control signals are based on the temporal derivative of controlled variable (DiStefano et al., 2012). We note that derivative-like responses of HONs to glucose (Figure 1) closely resemble such signals (DiStefano et al., 2012). On the other hand, HON subsets that signal absolute glucose fluctuations (Figure 5) resemble proportional sensors in engineering (DiStefano et al., 2012). Closed-loop control is more effective when it uses derivative sensors, because their anticipatory-like responses preempt big deviations from a set-point (DiStefano et al., 2012). In HONs, the derivative-based output may shift the body to energy storage based on how fast the glucose is rising, while the proportional output may fine-tune physiology according to actual blood glucose. Finally, multiplexing locomotion information and these glucose-dependent signals in individual HONs may more efficiently utilize the limited resources of the brain for neural coding (Akam and Kullmann, 2014; Panzeri et al., 2010).

In summary, our findings identify the HON system as a neural substrate for representing proportional and derivative blood glucose dynamics in the brain, and as an essential mediator of glucose-induced behavior. Deeper knowledge of how precisely-defined neural classes and connections turn blood glucose into adaptive responses will facilitate insights into blood glucose-associated behavioral changes in health and disease (DeFronzo et al., 2015; Gailliot and Baumeister, 2007; Gold, 1995; Messier and Gagnon, 1996; Zhou et al., 2020).

## METHODS

### Experimental subjects

All animal experiments followed United Kingdom Home Office regulations or Swiss Federal Food Safety and Veterinary Office Welfare Ordinance (TSchV 455.1, approved by the Zurich Cantonal Veterinary Office). Adult C57BL6 male mice were studied, thus whether conclusions apply to female mice remains to be determined. For HON deletion experiments (Figure 5 and Supplementary Figure 5), we the previously validated orexin-DTR mice (Gonzalez et al., 2016b; Viskaitis et al., 2022); confirmatory histology shown in Figure 5A was performed as in (Viskaitis et al., 2022). Animals were housed in reversed light-dark cycle (lights off at 7:00 a.m.), and all experiments were performed during the dark phase. Animals had *ad libitum* access to food (3430 Maintenance Standard diet, Kliba-Nafag) and water, unless stated otherwise. Known effect sizes and variation were used for power calculations and determination of required number of animals, where possible, to maximize chances of meaningful results without using excessive numbers of experimental animals. Studies were repeated in at least 2 independent cohorts and used a semi-randomized crossover design.

### Surgeries and viral vectors

HON activity measurement or optogenetic control was achieved using orexin promoter (hORX)-driven GCaMP6s sensor or ChrimsonR excitatory optogenetic actuator, respectively; the specificities of these constructs for hypocretin/orexin neurons was previously validated by histological analyses (Duffet et al., 2022; Gonzalez et al., 2016b; Karnani et al., 2020). Briefly, the GCaMP6s calcium indicator was delivered using AAV1-hORX-GCaMP6s.hGH (10^13-10^14GC/ml, Vigene Biosciences). For fiber photometry studies, the GCaMP6s AAV was stereotaxically injected into the LH bilaterally (anteroposterior [AP], −1.35; mediolateral [ML], ± 0.90; dorsoventral [DV], −5.70, −5.40 & −5.10, 70 nl per site) and optic fiber cannulae (200 μm diameter, 0.39 numerical aperture (NA) fiber with 1.25 mm ceramic ferrule; Thorlabs) were implanted above the LH (AP, −1.35; ML, ± 0.90; DV, −5.00). For 2-photon imaging, the same virus and coordinates were used, but surgery was performed only on the left hemisphere and a GRIN lens (0.39 NA, 7.3 mm long, 0.6 mm, Inscopix) was slowly (150 μm/min) implanted instead of the cannulae. Implants and a custom-made aluminum head plate were secured to the skull using three skull screws and dental cement (Kemdent and C&B-Metabond). For optogenetic experiments, surgeries were performed in the same way as for the photometry studies, but delivering ChrimsonR using AAV9-hORX-ChrimsonR-mCherry (2×10^12 GC/ml, UZH Viral Vector Facility).

For intragastric infusion experiments, the surgery was adapted from rats as previously described (Viskaitis et al., 2022). A custom-made catheter was implanted by guiding tubing subcutaneously through the dorsal to the ventral incision. A fixation mesh was then positioned subcutaneously at the level of the scapula blades and catheter exited on the animal’s back. To prevent damage, a custom-made cover was placed on the catheter after every experiment. To prevent catheter blockage, saline (~0.1 ml) was used to flush catheters daily for 5 days after surgery, and then on every second day until the end of the study. Catheter functionality was confirmed by the presence of back-flow after saline IG infusion and terminally by dissection. In the event of catheter blockage, subjects were excluded from further experiments, but the data collected prior to blockage were included in the analyses.

Glucose and temperature telemeters (DSI) were implanted following the manufacturer’s instructions. The glucose sensor entered the systemic blood system via the left carotid artery with ~1.5 mm of the sensor protruding into the aorta. All surgeries were performed under aseptic conditions and animals received isoflurane anesthesia, as well as operative and postoperative analgesia.

### Fiber photometry

For fiber photometry experiments (Figures 1,2), we used a custom camera-based photometry system (built with assistance of Dr. Dale Elgar, INSS and based on (Kim et al., 2016)). Alternating illumination of two excitation LEDs (405 nm and 465 nm at 20 Hz each, average power =100 μW at implant fiber tip) was used to record LH HON-GCaMP6s emission fluorescence bilaterally, in 1-3 animals simultaneously. Emission generated by the 405 nm LED was used as a control for movement artefacts, the effects of which were further minimized by recording in habituated, head-fixed animals (Figure 1A). GCaMP6s bleaching, fiber illumination or expression variability were accounted for by de-trending and normalizing each trace as follows: 1) local minima of each trace were found (“convhull” function in Matlab); 2) a least-squares triple exponential was fit through the convex hull; 3) each trace was detrended by subtracting and dividing by its minima-fit; 4) each trace was z-score normalised based on its 20 min pre-infusion standard deviation (SD) and mean.

### 2-photon imaging and data analysis

Volumetric 2-photon imaging via GRIN lenses was performed as previously described (Karnani et al., 2020). Excitation of GCaMP6s was achieved with a femtosecond-pulsed mode-locked Ti:sapphire laser (Spectra-physics Mai Tai HP Deepsee 2) at 950 nm. The emission fluorescence was imaged using a resonant/galvanometer scan head two-photon microscope (INSS), equipped with a 20 × (0.45 NA, Olympus) air-IR objective, a custom electro-tunable lens and a 510/80 nm band-pass emission filter. A volume of 512 × 512 pixels x 6 planes was recorded at 5.1 volumes/s using a custom Labview software. Resultant image stacks were processed in FIJI, Matlab, and Python softwares as follows: 1) the imaged volume was separated into separate planes (and lens-transition plane was discarded from further analysis); 2) 2×2 binning and TurboReg (precise, rigid) motion correction were applied; 3) cell outlines were manually drawn and labelled as regions of interest (ROIs); 4) ROI maps were applied across sessions within the same animal and adjusted; 5) ROIs that corresponded to the same cell in a plane or across neighboring planes were identified (in at least 2 independent experimental sessions, >5% ROI overlap of 3 pixel expanded contours, > 90% cross-correlation coefficient and <2s lag) and joined in further analysis; 6) HON cell activity was aligned based on saline/glucose infusion timing and same condition recordings were averaged per each cell individually. 7) HONs were assigned to glucose-response classes by fitting their activity profile to transformations of glucose dynamics (inverted-sign derivative, derivative, proportional, and inversely proportional); 8) classified cells were anatomically mapped.

For classification of HON activity to transformed glucose dynamics, templates were constructed using the average blood-glucose trace following IP-injection of 2 g/kg glucose from mice not used in the classification. The mean trace was smoothed using a 1.5-minute window moving mean before relevant transformations were applied. If the transformation was a derivative, the template was smoothed again after differentiation using the same filter parameters. Cell activity traces were extracted and aligned to the first 20 minutes of the template-trace following glucose IP injection. By calculating the Pearson’s Correlation Coefficient (PCC) of the extracted traces with each of these templates, we assigned cells to the response-class with the maximum PCC. A two-sided p-value was calculated alongside each correlation (SciPy library). Using the Bonferroni-adjustment for multiple comparisons, we classified responses that had no p-value with any template less than alpha = 0.001/4 as belonging to a fifth‘no-response’ category.

HONs were classified with regards to their correlation to running, by correlating whole session activity trace with locomotion. Positively correlated (p<0.05, rho > 0.01), negatively correlated (p<0.05, rho < −0.01) and noncorrelated cells (p>0.05 or rho between −0.01 and 0.01) were identified by using Spearman’s correlation.

### Concurrent monitoring of glucose, metabolic parameters, and locomotion

Blood glucose concentration and flank body temperature were measured and pre-processed with the HD-XG telemetry system (DSI). Metabolic measurements were obtained by recording continuous gas exchange in a custom enclosure by an adapted Field Metabolic System (Sable). Running of head-fixed animals was measured on a wheel using an optical encoder (Honeywell, 128 ppr 300rpm Axial). State changes of the encoder were recorded using a master Matlab code running photometry or Labview programs synced to the 2-photon microscope. Metabolic data was pre-processed in ExpeData software, z-transformed to account for sensor lag, and exported for analysis in Matlab. Respiratory energy expenditure was calculated using Weir’s formula: EE (kcal/min) = (3.94 x VO2(l/min)) + (1.1 x VCO2(l/min)). All acquisition systems were synced via digital signals to the Matlab code running photometry or to the Labview 2P imaging program. All data was exported and further processed in Matlab. Simultaneous measurements were resampled to achieve the same acquisition rate of 1Hz. For visual clarity, glucose and photometry data in Figures 1 and 2 were smoothed with a 10 minute moving mean and down-sampled to 20 seconds. To generate hysteresis plots of HON activity vs blood glucose from diverse infused glucose parameters Figure 1, boundaries of the blood glucose peak were identified and 50 equally spaced points were taken to plot HON signal vs blood glucose level (Figure 1H) or vs blood glucose derivative (Figure 1K). Classic glucose tolerance tests (Supplementary Figure 5A,B) were performed as in our previous work (Gonzalez et al., 2016b).

### Glucose doses

A critical feature of our experimental design is that, because of the rapid physiological counter-regulation of glucose in the body, in all our analyses of HON population responses to glucose we used actual measured blood glucose values, rather than the administered doses of glucose. In this way, we assessed the effects on neural activity of a large range of blood glucose concentrations (baseline range = 3.5-11.1 mM; baseline mean = 7.3±0.17 mM; glucose peak range = 11.1-34.9 mM, mean = 24±0.9 mM) and rates of change (range = 0.12-1.45 mM/min, mean of maximum rate of change = 0.56±0.03 mM/min). To achieve this range of glucose variation, our glucose infusions varied in both concentration and route of administration (intragastric, IG; or intraperitoneal, IP). Specifically, for IG infusions, we used a range of concentrations (0.08, 0.12, 0.146, 0.24 and 0.45 g/ml), infusion rates (50, 66.7, 90 and 180 μl/min), and volumes (<0.45 ml for fast 180 μl/min infusions and <0.9 for the other rates). For IP glucose infusions, a 2 g/kg dose was achieved by injecting 100-150 μl. All infusion parameters were used to generate data analyzed in Figure 1. The 0.24 mg/ml IG dose (delivered at 90 μl/min rate over 10 minutes) was used in Figure 2, and in Supplementary Figure 2 the findings are confirmed for a wider range of infusion parameters (as indicated in the figure legend of Supplementary Figure 2). 2g/kg IP dose was used in Figures 3–5. Note that the routes of administrations and doses are provided here for the sake of completeness, since our study relates blood glucose (directly measured, rather than inferred from injected doses) to HONs activity, and in relationship between blood glucose and HON activity is similar across different methods of glucose injection (Supplementary Figure 1, IP and IG compared in figure legend).

### Encoding model

To determine relative contributions of various behavioral and metabolic variables to HON responses, we used a generalized linear model approach, based on principles described before (Engelhard et al., 2019). For this model, HON population activity was used as the response variable, while predictor variables were running, blood glucose, body temperature, consumed O_2_ volume, produced CO_2_ volume and their derivatives with respect to time (Figure 2H). All variables were down-sampled to 1 min in order to equalize their sampling rates and focus on the slow dynamics, Savitzky–Golay filtered (1st order, 5 sample window), and normalized by z-scoring. Resultant data was fit using “glmfit” function in Matlab by bootstrapping ¼ experiment-duration chunks randomly over 2000 iterations. Here 70% of data was used for training and 30% was used for validation on untrained data. Non-bootstrapping methods such as leave-one-out cross validation were also used to confirm the findings. For each fitting iteration, partial models based on the same training data but without a single independent variable were generated. Then a coefficient of determination (R2) was calculated for full and partial models on the validation data (either on the 30% of bootstrapped data or the left out experiment). We determined relative contribution of a given predictor to HON activity dynamics by comparing how much the encoding model performance has declined without a given variable - by comparing the R2 of the partial model to the R2 of the full model (Figure 2I). Negative relative contributions were set to zero, following the method described in (Engelhard et al., 2019).

### Behavioral analysis

HON-ablated and their respective control mice were produced by injecting diphtheria toxin (Sigma D0564; 1 mg/ml; 0.1 ml) via intraperitoneal (IP) route into both DTR+ and DTR-mice, as previously described (Gonzalez et al., 2016b). Constitutively active HON state was produced by optogenetic LH stimulation of HON-Chrimson mice, achieved by tonic 10 Hz, 5ms pulses of 635 nm laser. In both cases, HON-manipulated and control mice were injected with either glucose or saline (in a randomized cross-over fashion over 2 consecutive days) and placed into an open-field arena (40×40 cm). Behavior was scored using video analysis of the first 10 minutes by Ethovision XT (Noldus) and by a custom-made machine learning classifier as described in detail previously (Viskaitis et al., 2022). The machine learning classifier was trained on ~1200 labeled examples to identify 5 separate behaviors: grooming, rearing, resting, running and turning or sniffing. Additional >300 labelled examples were used to train a behavioral classifier for optogenetic-cable tethered mice. Separate running bouts were defined as occurrences of movement above a threshold (18cm/s for untethered or 10 cm/s for tethered mice) for at least 1 s/bout, separated by more than 2 s. Similarly, when machine learning was used to identify forward locomotion directly, separate running bouts were identified when spaced by at least 2 s.

### Data analysis and statistics

Raw data were processed in Matlab. Statistical analysis was done in GraphPad Prism 9.0, Matlab or Python. Key comparisons between saline and glucose, and cell classification analyses, were performed on raw, non-smoothed data. Sample size, statistical tests used and their results are indicated in the figures, their legends and/or their descriptions in the text. Statistical comparisons were performed on non-filtered data, but some traces were filtered for visual purposes, as indicated. Statistical analysis was based on settings recommended by GraphPad Prism 9.0. P values of <0.05 were considered significant. Where significance is presented, p values are as follows: *p<0.05, **p<0.01, ***p<0.001, and ****p<0.0001, ns = p>0.05. Outliers that failed a ROUT 1% test were removed. Data are presented as means and standard error of the mean (SEM) unless stated otherwise.

---

**Supplementary Figure 1.**
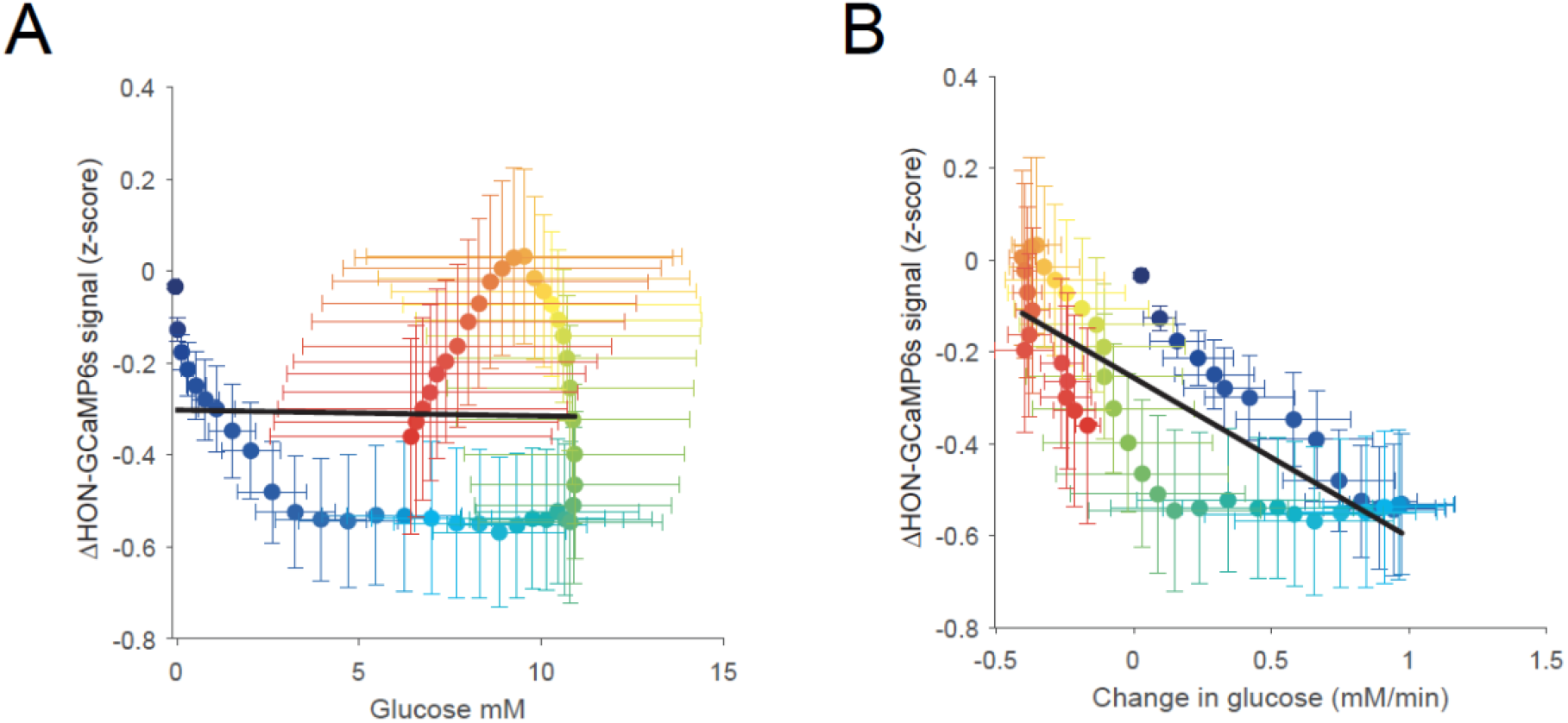
Relationship between blood glucose and HON activity after IP injection of glucose. Related to Figure 1. (A) HON activity did not have a linear relationship with varying blood glucose after intraperitioneal (IP) injection of glucose. Linear fit did not explain variability and fit slope was not different from zero; R2 = 0.0005, p =0.87. (B) There was a significant linear relationship of HON activity with changes in blood glucose and the slope was significantly non-zero: R2 =0.64, p <0.0001. HON activity response slopes vs changes in blood glucose were not affected by the route of glucose administration: IG vs IP in Figure 2K vs Figure S2D; F test (1,96) 1.888=; p = 0.17. IP glucose responses quantified using 19 HON responses from 8 animals & 4 glucose recordings in 4 animals. Data presented as means and SEM.

**Supplementary Figure 2.**
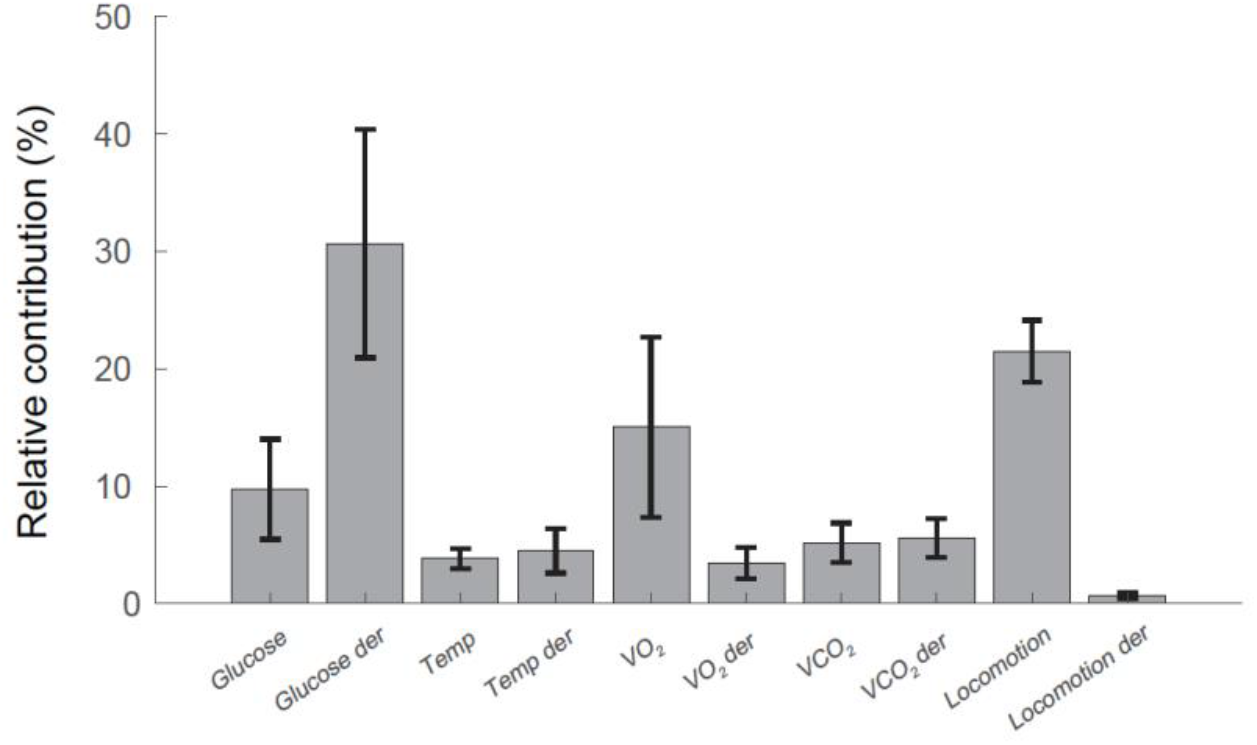
Glucose derivative is a top predictor across a wide range of glucose infusion parameters. Related to Figure 2. Unlike the dataset than shown in Figure 2 (0.24 g/ml, 90 μl/min for 10 min), here a wider range of the infused glucose parameters were used: glucose concentration was either 0.12, 0.146 or 0.45 g/ml, the infusion rate was varied between 66.7, 90 and 180 μl/min, and duration was either 2.5, 10 or 15 min. 11 sessions from 5 animals. Data presented as means and SEM.

**Supplementary Figure 3.**
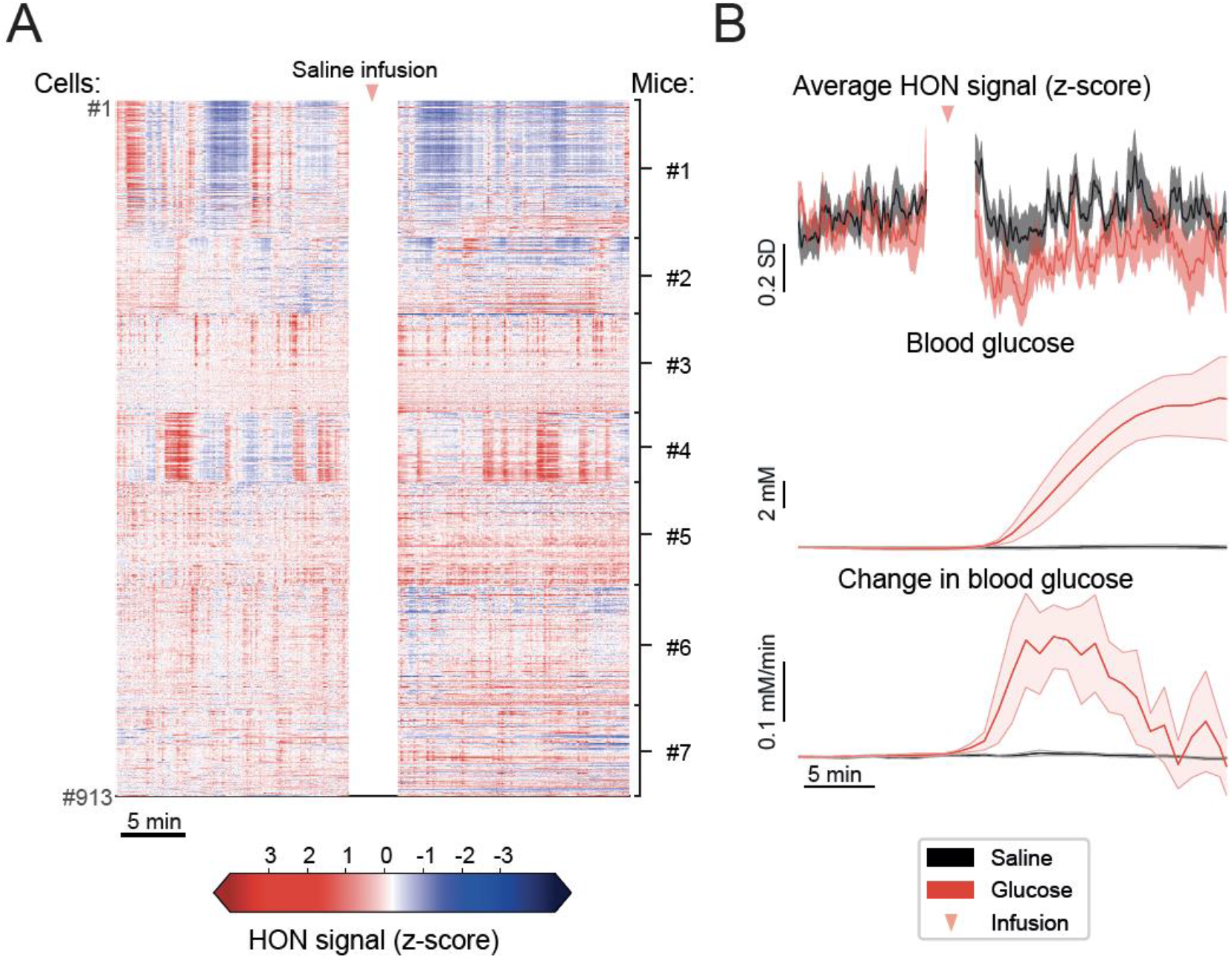
Single cell HON responses to saline and glucose infusions. Related to Figure 3. (A) Individual responses of 913 HONs to saline infusion from 7 mice. (B) Temporal alignment of average HON responses (top) to blood glucose concentration (middle) and its derivative (bottom) after saline or glucose infusions. Blood glucose data is from the same recording sessions as shown in SI. Breaks in the neural recordings are due to laser shutter being closed during infusions in majority of the experiments. Data are presented as means and SEM.

**Supplementary Figure 4.**
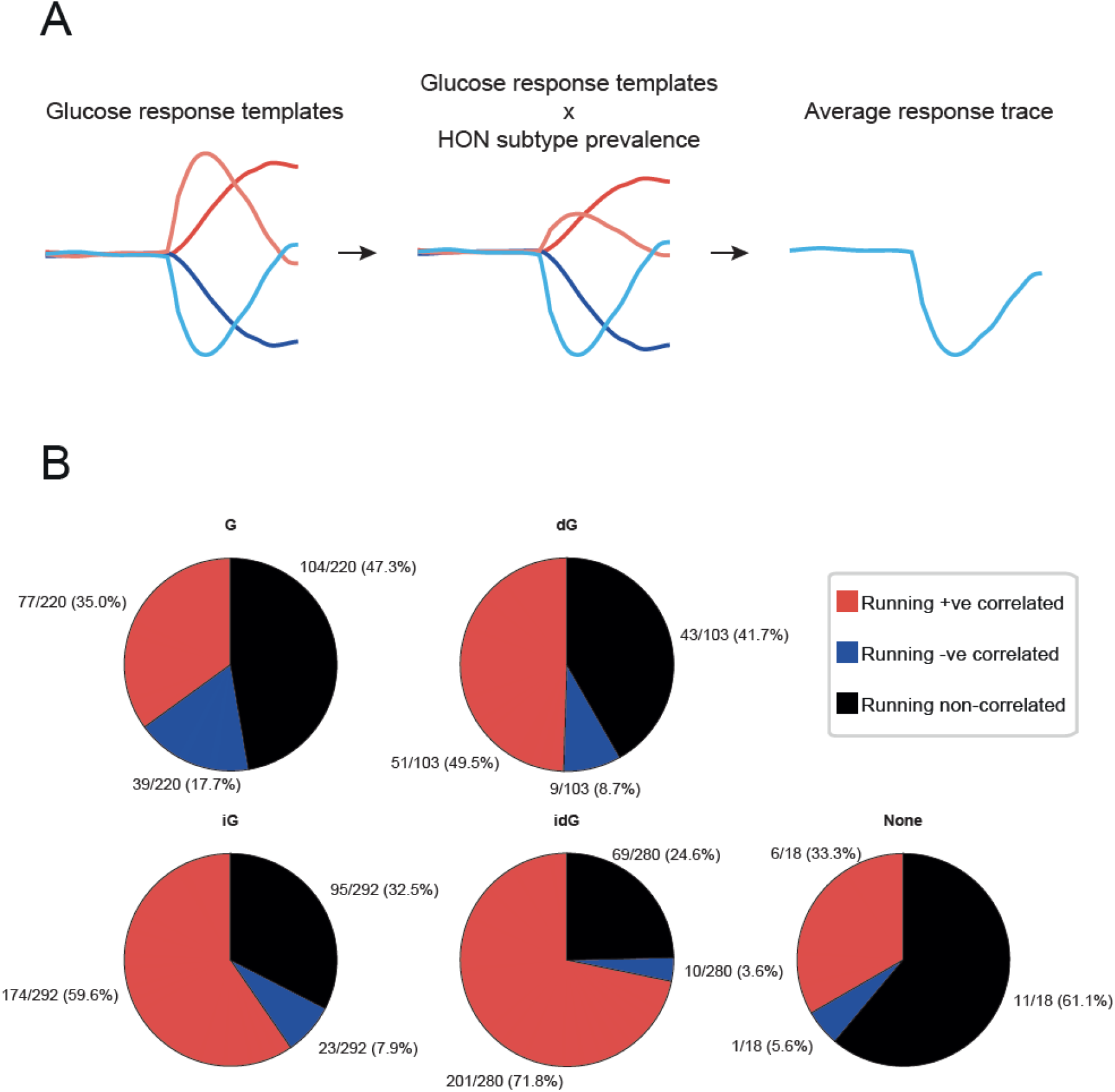
HON population response to glucose and correlation of single cell HON responses to glucose infusions versus running. Related to Figure 4. (A) Using blood glucose templates from Figure 4A and multiplying them by the relative prevalence of individual HONs that fit these profiles, we obtained an average response trace that resembles that of the actual HON population response after the infusion of glucose. (B) Glucose inhibited cells (iG and idG) tended to also have a larger proportion of +ve correlated running cells.

**Supplementary Figure 5.**
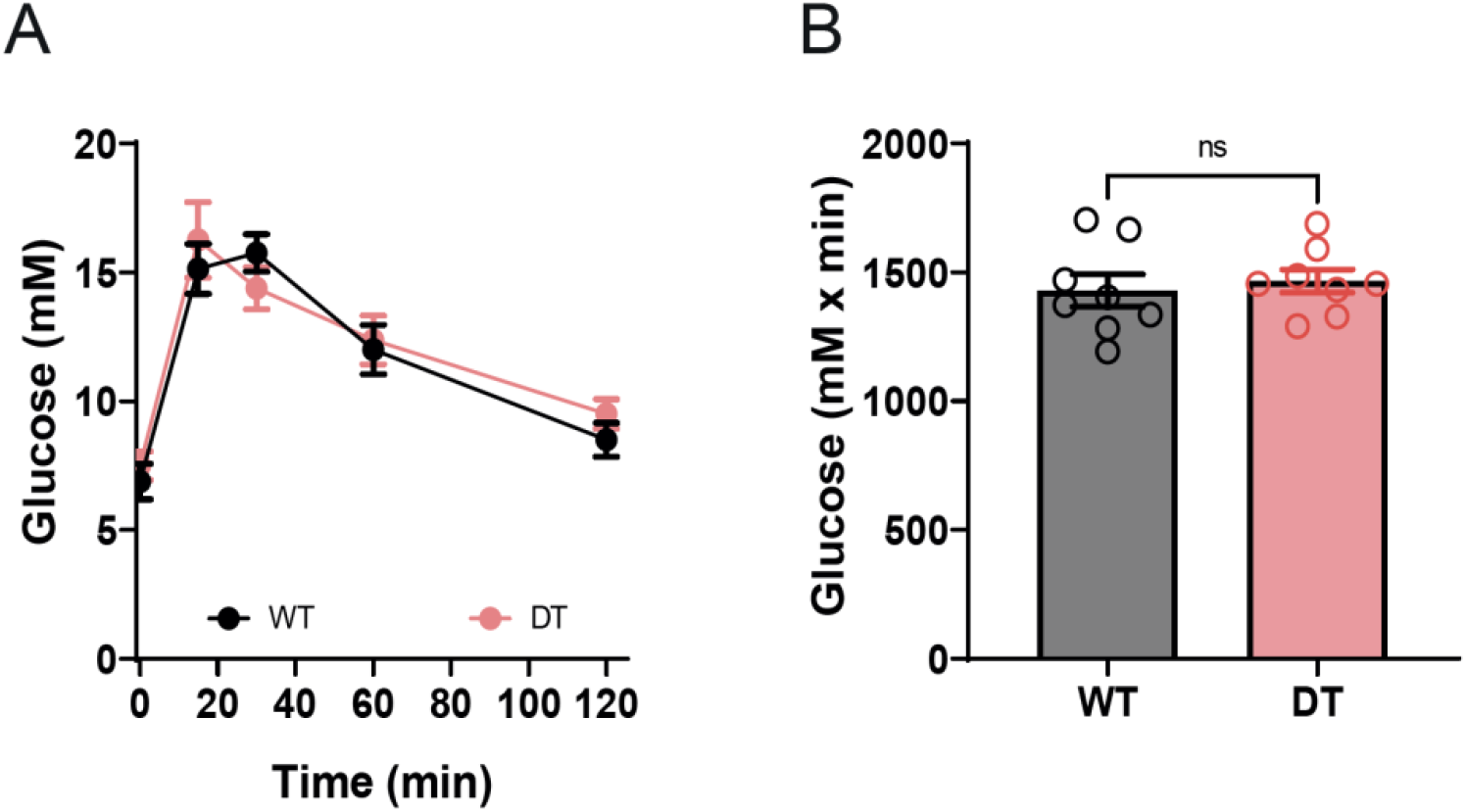
Glucose tolerance test (GTT) in wild-type and HON-ablated mice. Related to Figure 5. (A) GTT timeline. (B) GTT area under the curve. Unpaired t-test p =0,56, t=0.59, df=l4. Data presented as means and SEM.

## ACKNOWLEDGEMENTS

This work was funded by ETH Zürich, and The Francis Crick Institute which receives its core funding from Cancer Research UK (FC001055), the UK Medical Research Council (FC001055), and the Wellcome Trust (FC001055).

